# SNP analysis implicates role of cytosine methylation in introducing consequential mutations in *Vibrio cholerae* genomes

**DOI:** 10.1101/070839

**Authors:** Mohak Sharda, Aswin Sai Narain Seshasayee, Supriya Khedkar

## Abstract

Epigenetic modifications play a key role in gene regulation and in recognition of self DNA in bacteria. In-spite of their positive role in cell survival, modifications like cytosine methylation incur a mutational cost. Cytosine methylation, specifically 5-methylcytosine, is prone to hydrolytic deamination which leads to C → T and G → A transitions. Here, we first study the abundance of mutagenic cytosine methylation target motifs and show that bacteria like *Vibrio cholerae* might use motif avoidance as a strategy to minimize the mutational effect of deamination of methylated cytosine. Second by performing SNP analysis on whole genome sequence data from *Vibrio cholerae* patient isolates we show a) high abundance of cytosine methylation-dependent mutations in the cytosine methylation target motif RCCGGY, b) 95% of these C → T and G → A transitions in the coding region lead to non-synonymous substitutions and c) many of these transitions are associated with membrane proteins and are implicated in virulence. Thus, our SNP analysis of *V. cholerae* genomes implicates the role of cytosine methylation in generating genotypic diversity with adaptive potential.

---

Cytosine methylation is as an epigenetic signal that modulates gene expression (Chao et al., 2015; Deaton and Bird, 2011; Kahramanoglou et al., 2012). In higher eukaryotes, many processes involved in organism development are regulated by methylation at CpG islands (Smith and Meissner, 2013). In bacteria however, the origins of DNA methylation might be traced to selfish elements coding for restriction-modification (R-M) systems, which contribute to defense against invading DNA such as phages (Kobayashi, 2001). R-M systems typically comprise a sequence-specific endonuclease activity (restriction), and a DNA methyltransferase activity that modifies DNA and protects it from cleavage by the restriction endonuclease (Tock and Dryden, 2005). R-M systems are harbored by both chromosomal and plasmid DNA, and the two components are generally present together as a pair. Loss of the gene encoding a restriction enzyme produces an orphan or solitary methyltransferase (Ershova et al., 2012; Seshasayee et al., 2012). Whereas the presence of an R-M system can select against the presence of sequence motifs recognized by the system for cleavage (Qian and Kussell, 2012; Rocha et al., 2001; Rusinov et al., 2015), the generation of a solitary methyltransferase can relax such selection. A few solitary DNA methyltransferases have been characterized, and these studies have indicated (a) roles for these in regulating the expression of several genes, including those involved in core processes such as the cell cycle (Gonzalez et al., 2014; Marinus, 1987; Messer et al., 1985; Reisenauer et al., 1999) and (b) their role in protecting the genome from attack by selfish restriction-modification systems (Takahashi et al., 2002). These are in addition to their well-known roles in helping initiate DNA replication and in mismatch repair (Pukkila et al., 1983; Reisenauer et al., 1999).

Though bacterial DNA methylation can target both cytosine and adenine bases, our emphasis here will be on the generation of 5-methyl cytosine (5mC). Despite the emerging positive roles for 5mC in regulating gene expression, it presents a cost. Cytosine deamination is a common source of spontaneous mutation, in which C is converted to U, which can be specifically recognized and corrected (Krokan et al., 2002). Deamination of 5mC occurs at a rate higher than that of C (Shen et al., 1994). Additionally, factors like single stranded DNA (Frederico et al., 1990), high temperature (Ehrlich et al., 1986) and flanking DNA (Fix and Glickman, 1986; Lutsenko and Bhagwat, 1999) significantly increase the rate of deamination. Deamination of cytosine also produces T, which being a natural component of DNA becomes more difficult to detect as a mutation, especially under situations in which base-pairing incompatibilities cannot be exploited for repair. For example, CpG islands, which are targets for the generation of 5mC in eukaryotes, contribute to about 30% of point mutations in the human genome (Cooper and Youssoufian, 1988). This is in-spite of the existence of demethylases (Bhattacharya et al., 1999; Detich et al., 2002) that can reverse methylation at CpGs, and repair mechanisms like base excision repair mechanisms that can revert C → T and G → A transitions (Walsh and Xu, 2006). Further, the depletion of CpG islands due to C → T and G → A transitions is implicated in various diseases and disorders (Cooper and Youssoufian, 1988), including cancer (Jones and Baylin, 2002; Walsh and Xu, 2006). However, a recent paper has indicated that higher mutation rates in CpG islands in Drosophila may not be due to the mutagenic potential of 5mC per se, but due to the presence of these sites in transcriptionally active regions, or by chromatin-associated constraints on transcriptionally inactive loci (Glastad et al., 2016). Thus, the role of DNA methylation in generating mutations and genetic diversity in eukaryotes remains debatable.

In bacteria, demethylases are absent and repair mechanisms like mismatch repair revert C → T and G → A transitions only during exponential growth (Jones et al., 1987; Zell and Fritz, 1987). In stationary phase, which might comprise a large part of the bacterial life cycle in the wild, the only reported repair mechanism is the *E. coli* Very Short patch Repair (VSR) which is encoded adjacent to the orphan DNA cytosine methyltransferase Dcm. VSR can repair C → T mutations within the CCWGG motif targeted by Dcm (Lieb, 1991; Macintyre et al., 1997). Some other bacterial species appear to lack such a repair mechanism. For example *Vibrio cholerae*, the causative agent of cholera, encodes an orphan cytosine methyltransferase (VchM), which methylates the outer cytosine in the RCCGGY motif (Banerjee, 2006), but lacks a VSR-like repair system. It has been demonstrated, using tests for antibiotic resistance, that wildtype *V. cholerae* display a higher mutation rate than a *vchM-* strain (Banerjee, 2006). However, the genomic landscape of 5mC → T mutations in *V. cholerae*, and the question of whether there might be indications of selection in favor of the deployment of DNA methylation as a source of generating increased protein sequence diversity on a genomic scale remains unanswered. We seek to answer this question here.

## Materials and methods

### Data

The reference genome sequence data for *E. coli* K12 MG1655 (NC_000913) and *Vibrio cholerae* El tor N16961 (NC _ 002505 – Chromosome 1 and NC _ 002506 – Chromosome 2), and ~300 other bacterial genomes used in this study was obtained from NCBI (Table S1). The whole genome sequence data for the five datasets namely the 7^th^ Pandemic, Bangladesh, Chandigarh, Nepal and Haiti were obtained from NCBI (accession numbers of these strains are listed in **File_S1**)

The ORF annotations for all the genomes used in this study was obtained from NCBI (.ffn) files. Information on the proteins and rRNA sequence data and gene positions was extracted from NCBI (.faa), (.frn) and (.ptt) files respectively.

The protein sequence of 5-methylcytosine methyltransferases, the methyltransferase target motifs and the organisms in which they were found was obtained from REBASE http://rebase.neb.com/cgi-bin/seqsget?L+ad‚ accessed in March 2016. Based on the organism name and strain information the NCBI accession numbers for all the organisms used in this study were extracted from the NCBI genome list. A list of all type II methyltransferases was obtained from http://rebase.neb.com/cgi-bin/seqsget?L+md and compared with the 5-methylcytosine methyltransferase list to obtain all the type II 5-methylcytosine methyltransferases.

### Annotation of orphan methyltransferases and *usr+* genomes

A methyltransferase was annotated as orphan if its homolog – as determined by phmmer search against individual bacterial protein sequences (NCBI .faa files) with an e-value cutoff of 10-^100^; was not detected in the vicinity (10 genes upstream or 10 genes downstream) of a restriction enzyme with same target motif as that of the methyltransferase (**Table S1**), as described earlier (Seshasayee et al., 2012).

Protein sequences of VSR (very short patch repair) proteins were obtained from uniprot. A multiple sequence alignment was generated for these 452 amino acid sequences using Muscle (Edgar, 2004) with default parameters. A profile hmm was generated for VSR protein and was searched against bacterial protein sequences (NCBI .faa files) to identify homologs of VSR proteins in the respective bacterial genomes (**Table S1**) using hmmbuild (default settings) and hmmsearch (default settings) commands respectively. This homolog search was implemented using HMMER (Finn et al., 2011). For bacteria encoding for multiple 5-methylcytosine methyltransferases, the identified VSR homolog was assigned to a particular methyltransferase depending on its proximity (10 genes upstream or 10 genes downstream) to the methyltransferase.

### Determination of motif counts, motif positioning and type of substitutions

The number of target motifs in the respective genomes was counted using the respective whole genome sequence information from the reference strain using an in-house python script. We also calculated expected counts for each motif in the respective genomes by performing 1000 iterations of genome shuffling. We computed fold change for motif counts by calculating the ratio of observed motif counts and average expected counts.

To count motifs in the protein coding region, ORFs from respective genomes were considered. For counting the number of motifs in the non-protein coding region the total number of motifs in the organism was subtracted from the number of motifs identified in the coding region.

We determined the percentage of non-synonymous substitutions at the 5-methylcytosine target motifs by considering whether a C → T transition at the methylated base leads to change in the amino acid for motifs present in the coding region. The distribution for the random expectation of percentage non-synonymous substitutions for all the C −> T transitions in the genome was obtained by performing 50 iterations of genome shuffling.

### Determination of enrichment of C->T mutations in the RCCGGY motif in *Vibrio cholerae*

We determined the total number of RCCGGY and XCXXXX from the reference genome of *V. cholerae.* The total number of C->T and G -> A transitions in each strain for each respective data-set was extracted from the output.vcf files obtained after running Breseq (Deatherage and Barrick, 2014). These transitions were segregated into three types 1) C->T transitions observed in the first C of the RCCGGY motif; 2) C->T transitions observed in the second C of the RCCGGY motif and 3) C->T transitions observed in the C anywhere except the first C of the RCCGGY motif (**Table_S2**). Enrichment for the C->T transitions in the first C within the RCCGGY motif was determined by comparing it to all other C->T transitions using Fisher’s exact test (**File_S2**). Similarly, the enrichment was also determined for the C -> T transitions at the first C as compared to the second C within the RCCGGY motif. The enrichment of G -> A transitions within the RCCGGY motif was computed in the same manner.

### Determination of Functional Enrichment

Functional annotations for individual genes with C ->T and G>A transitions were extracted from NCBI(.ptt files). These Cluster of Orthologous Genes (COG) annotations for the 70 genes with C>T and G>A transitions were used to determine enrichment of specific functions. 23 COG classes belonging to three broad functions – information storage and processing, cellular processes and signaling, and metabolism-were considered. A Fisher’s exact test and bonferroni corrected p-values were used to determine statistical significance for the enrichment of each COG class.

A (Gene Ontology) GO: annotation based enrichment analysis was also performed using KOBAS 2.0 (Xie et al., 2011) using Fisher’s exact test and Benjamini Hochberg 1995 FDR correction method.

### Phylogenetic analysis

16S rDNA sequence was obtained for 318 organisms used in this study. For organisms with multiple copies of rRDNA; one copy was selected at random and a multiple sequence alignment was generated by implementing Muscle (Edgar, 2004) in MEGA (Tamura et al., 2013) using default options. The generated alignment was used to construct a Neighbour-Joining tree using default settings in MEGA (Tamura et al., 2013). The information on the number of methyltransferases, the number of orphans and the presence of VSR was overlaid on the phylogenetic tree using iTOL(Letunic and Bork, 2016).

## Results

### Abundance of cytosine methyltransferase target motifs

Towards studying the extent and abundance of potential cytosine methylation-dependent mutations in bacteria, we counted the number of cytosine methyltransferase target motifs in two extensively studied model bacteria, *V. cholerae* and *E. coli*, that code for orphan cytosine methyltransferases VchM and Dcm respectively. We found that the VchM target motif RCCGGY is under-represented by about 30% (observed 2172; expected 3197) in the *V. cholerae* genome (**Figure 1A**). About a sixth of other proteobacterial genomes also showed avoidance of the RCCGGY motif in their genomes. On the other hand, in the *E. coli* K12 MG1655 genome the Dcm target motif CCWGG is over-represented by about 26% (observed 12050; expected 9497) (**Figure 1B**), and a similar trend was observed in other strains of *E. coli. E. coli* encodes for a VSR (very short patch repair) protein that reverts cytosine methylation-dependent mutations in stationary phase, this repair system is absent in *V. cholerae*. It has been hypothesized that the increased frequency of the CCWGG motif in *E. coli* might be a consequence of VSR erroneously converting CTWGG motifs to CCWGG (Bhagwat and McClelland, 1992). This analysis suggests that motif avoidance might be an alternative strategy used by bacteria like *V. cholerae* which do not encode VSR, to minimize the cost associated with cytosine methylation-dependent mutations.

**Figure 1:**
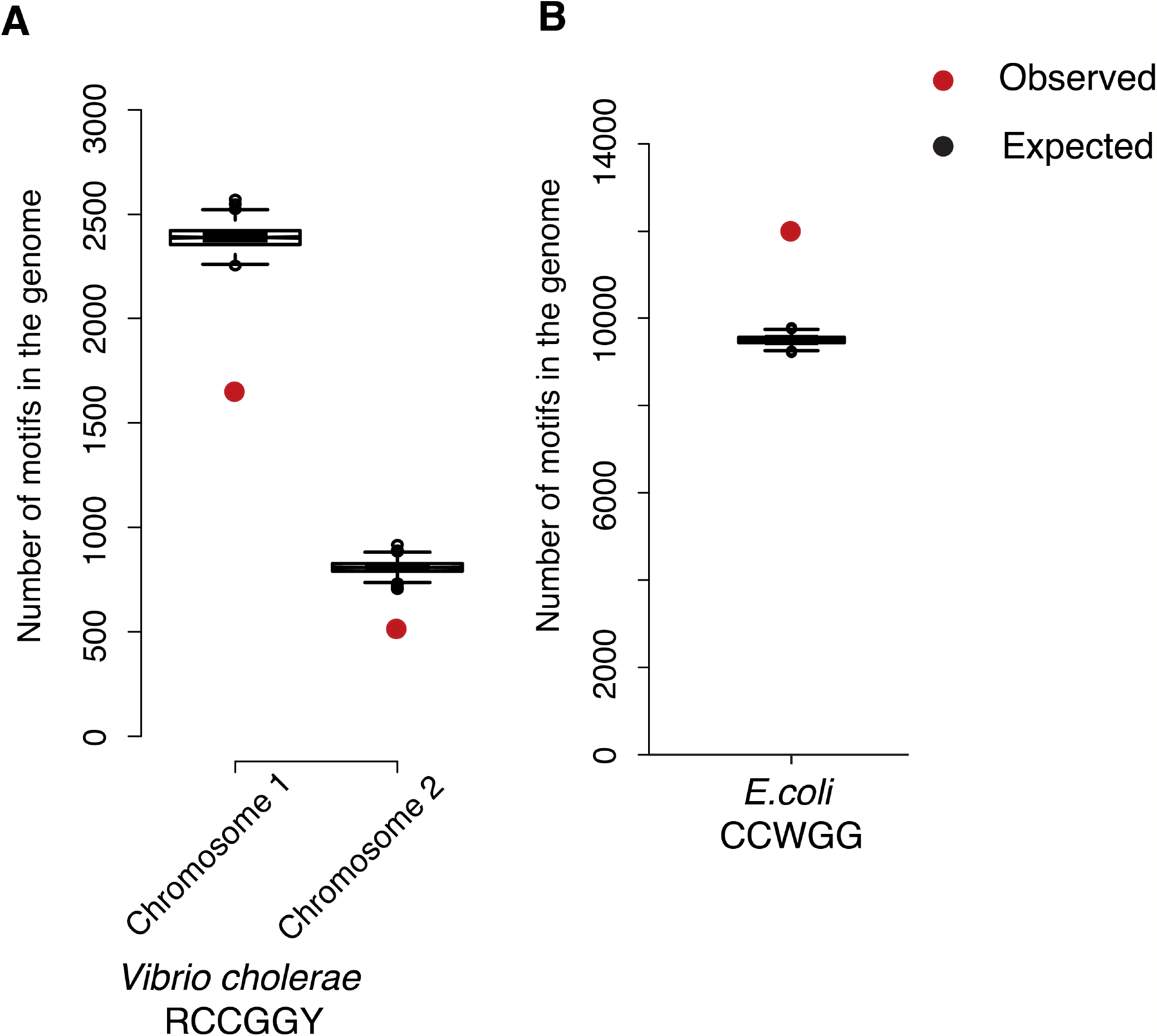
Abundance of *E. coli* and *V. cholerae* cytosine methyltransferase target motifs. **A.** Plot representing the under-representation of RCCGGY motifs in the *V. cholerae* genome in red, for both the chromosomes one and two. The expected motif numbers are obtained after performing 1000 random iterations of genome shuffling and are represented by the boxplots in black; **B.** Plot showing the observed over-representation of CCWGG motifs in the *E. coli* genome represented in red in comparison to expected motif counts obtained after 1000 random iterations of genome shuffling represented by boxplots in black.

Next, we determined the occurrence of the cytosine methyltransferase target motifs in the coding and non-coding regions of the respective bacterial genomes. We found that ~93% of the RCCGGY motifs in the *V. cholerae* genome and ~90% of the *E. coli* CCWGG motifs were encoded in the protein coding regions. We had previously noted that as many as 97% of methylated cytosines in coding regions are located in non-synonymous sites in the *V. cholerae* genome (Kahramanoglou et al., 2012). One would expect around 60% of any coding region base to be located in a non-synonymous site by random chance. Across our collection of type II 5-methylcytosine methyltransferases found in ~300 genomes, we find that on an average ~60% of 5mC positions are in non-synonymous positions; the distribution however is wide (**Figure S1**). Thus, taken together we suggest that in *V. cholerae*, there might be selection in favor of positioning 5mC at non-synonymous sites and that it might even be an exceptional case among other bacterial genomes.

### Mutational spectrum of methylated cytosines in virulent Vibrio cholerae El tor strains

The predictions about high abundance of non-synonymous substitutions at the RCCGGY motif in the *V. cholerae* led us to study the actual occurrence of C → T and G → A substitutions in sequenced *V. cholerae* genomes. Towards this end, we obtained whole genome sequence data for *V. cholerae* strains implicated in cholera epidemics from the early 1970s to 2013. The whole genome sequence information obtained from 4 different geographical locations and 3 different time periods are described in **Figure 2A and Table 1**. All the datasets comprise isolates of *Vibrio cholerae* El tor N16961 strains; this strain first emerged during the 7^th^ cholera pandemic. In total, for 287 temporally and geographically distinct *V. cholerae* El tor N16961 strains we identified single nucleotide polymorphisms (SNPs) against the fully sequenced reference genome of *V. cholerae* El tor N16961 (**Figure 2B**). We confirmed the presence of vchM in all these 287 *Vibrio cholerae* isolates.

**Figure 2:**
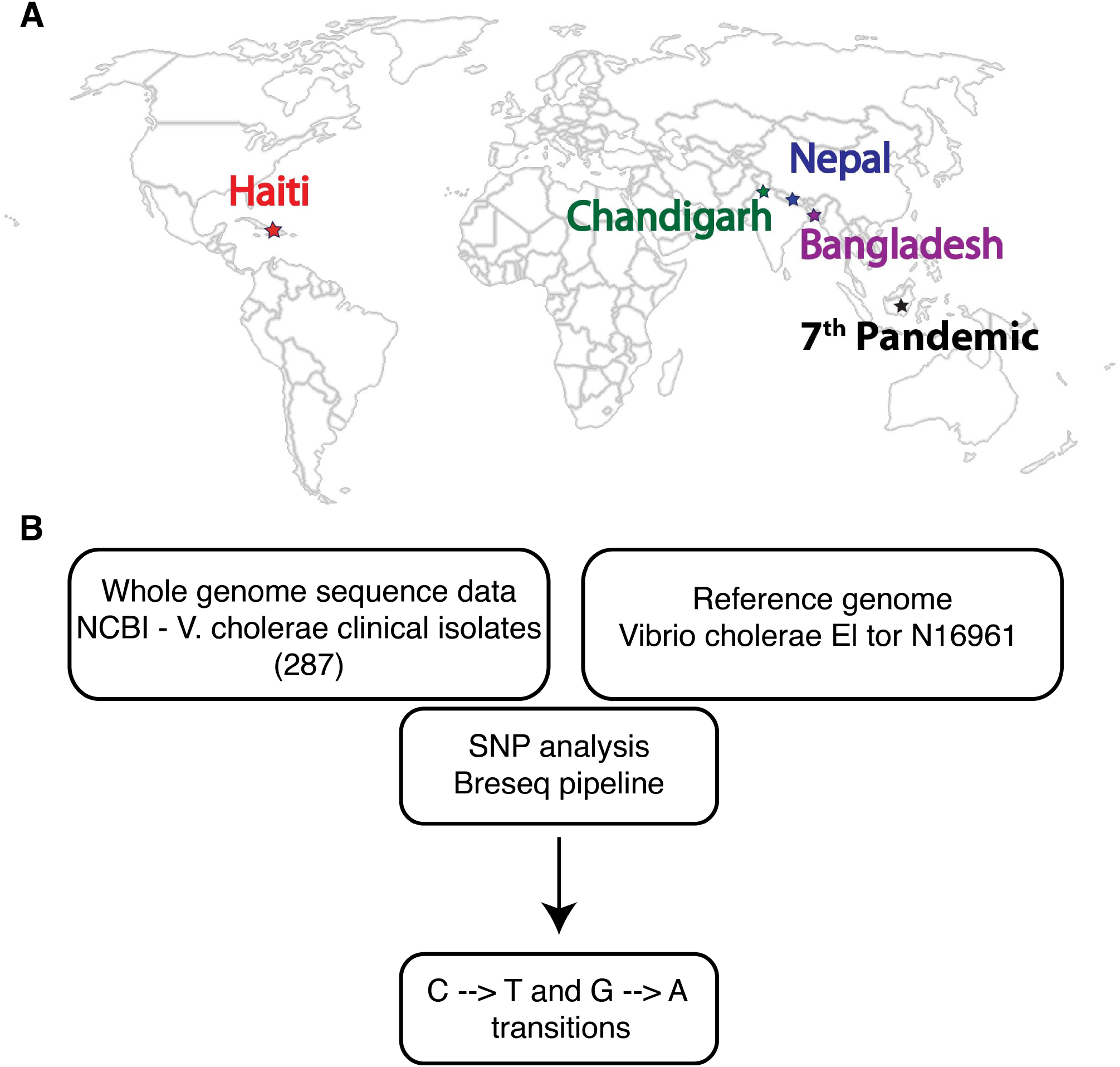
Datasets and analysis pipeline. **A.** World map highlighting the geographical distribution of the 5 datasets used in this study namely the origin of 7th Pandemic (black), Bangladesh (purple), Chandigarh (green), Nepal (blue) and Haiti (red); **B.** Flowchart depicting the analysis pipeline used to extract information on 5-methylcytosine deamination dependent transitions.

**Table 1.**

| Location | Year | Number of isolates |
| --- | --- | --- |
| 7<sup>th</sup> Pandemic | 1961-75s | 156 |
| Bangladesh | 2001-2011 | 50 |
| Chandigarh | 2009 | 38 |
| Nepal | 2010 | 23 |
| Haiti | 2013 | 10 |

To test the association between cytosine methylation and C → T and G → A transitions, we divided all observed C → T and G → A transitions into those that were within the RCCGGY motif and those that were outside. We first found that C → T and G → A transitions occurred more frequently within the RCCGGY motif than outside (**Figure 3A**). Next, to account for local differences in mutation rates, we compared the frequency of these transitions between the first and the second cytosine within RCCGGY. Note here that it is the first cytosine which is methylated. We found that the C → T and G → A transitions are significantly more frequent at the first cytosine when compared with the second (**Figure 3B**).

**Figure 3:**
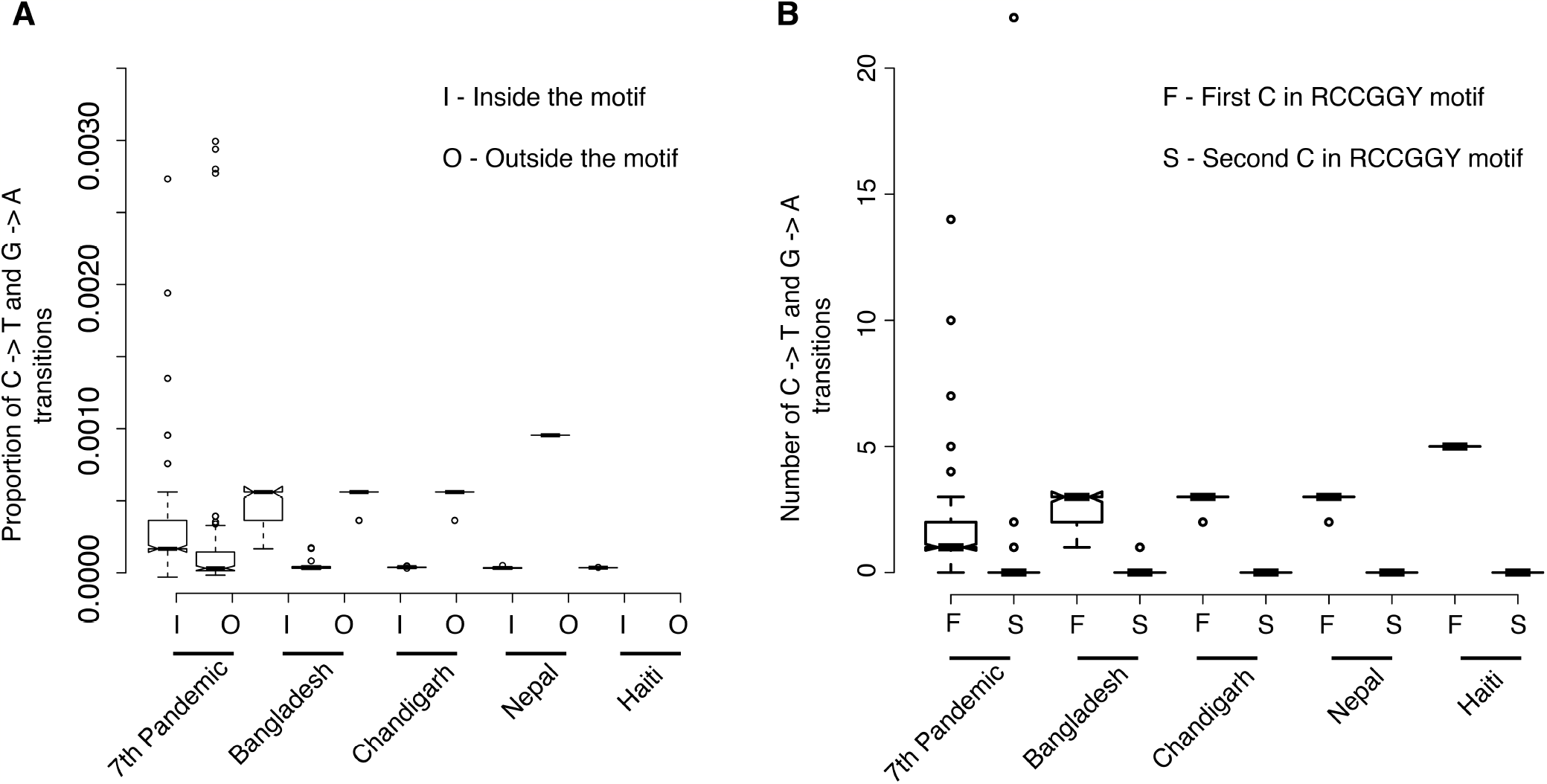
High C → T and G → A transition frequencies inside the VchM target motif RCCGGY **A.** Plot showing significantly high proportion of C → T and G → A transitions inside the *V. cholerae* RCCGGY motif as compared to outside the motif for the five datasets labeled in the figure (**File_S1**); **B.** Plot representing high proportion of C → T and G → A transitions corresponding to the first (methylated) C of the RCCGGY motif as compared to the second C (**File_S1**).

Further, we found that of the 2172 RCCGGY motifs in the reference *V. cholerae* genome, 81 showed a mutation in at least one of the 287 isolates considered in this study. On an average up-to 4 transitions were contributed by each of the pandemics. Many transitions were observed amongst the 7^th^ pandemic dataset from 1970s (**Figure 4 and Figure S2**). While 2 mutations were shared among all the 5 datasets, each dataset showed some degree of overlap with others (**Figure 4**).

**Figure 4:**
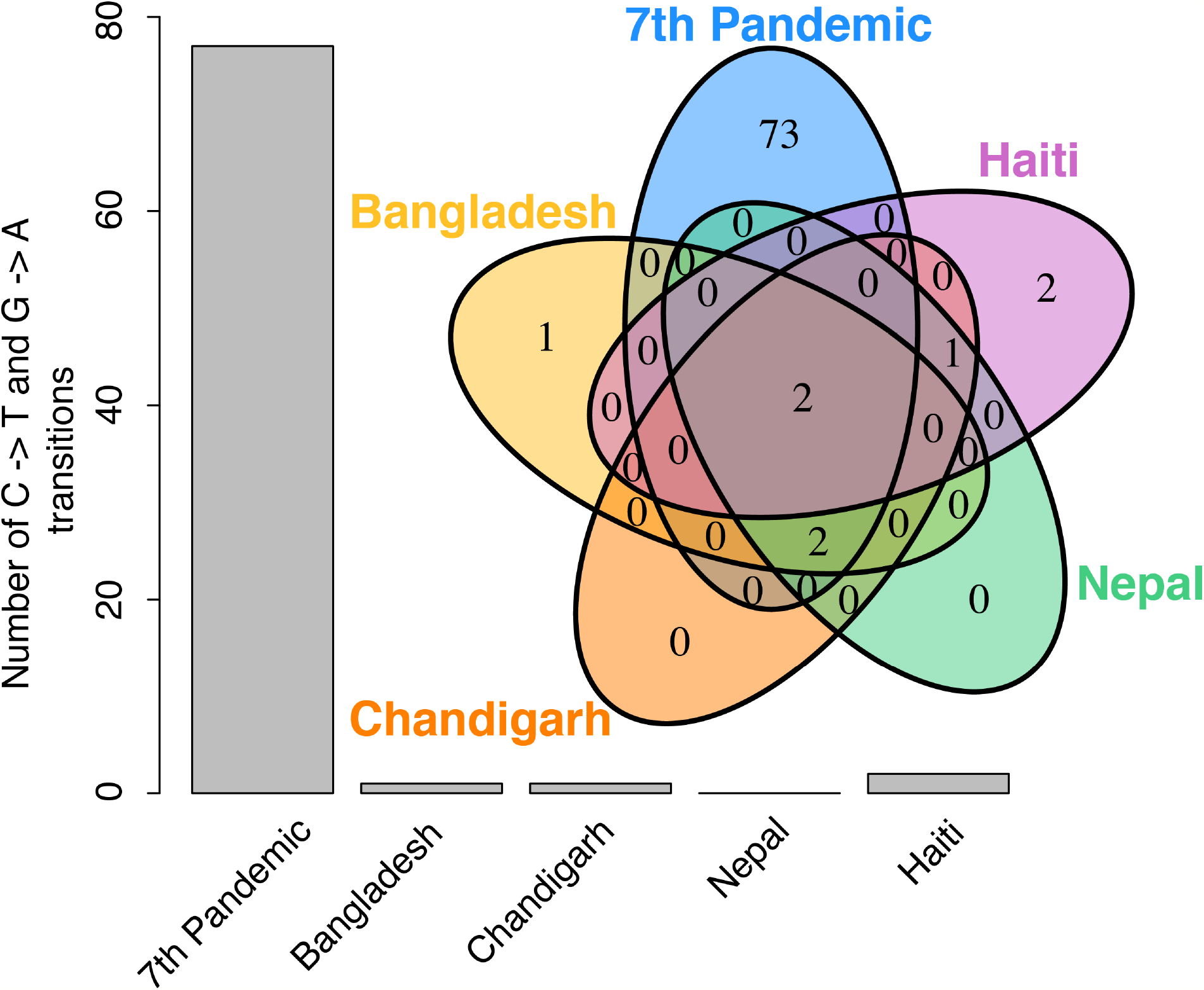
Plot representing the proportion of C → T and G → A transitions within the RC*CGG*Y motif in the five labeled datasets used in this study. Venn diagram represents shared C → T and G → A transitions among the marked datasets.

Of the total 81 C → T and G → A transitions, ~4 were found in the non-coding region and rest 77 were found in the coding region (**Figure S3**). ~95% (73 of the 77) of the C → T and G → A transitions in the coding region were at non-synonymous site as compared to random expectation of 66% for occurrence of non-synonymous C → T and G → A substitutions outside the motif.

These observations together implicate cytosine methylation in generating increased non-synonymous mutation rate at certain loci in *V. cholerae*.

### Functional consequences of cytosine methylation-dependent mutations

Non-synonymous mutations are often considered detrimental and are negatively selected for due to their effects on protein folding and stability (Pál et al., 2006). Hence, the presence of a significantly large proportion (95%) of methylation-dependent C → T and G → A at non-synonymous substitutions in *V. cholerae* patient isolates is rather surprising. To study this further, we extracted information on the functions of genes carrying these transitions (**Table S3**). We found that only 4% of these genes were horizontally acquired and that they show no functional enrichment for any COG (Cluster of Orthologous Genes) category. Further, we annotated the 77 genes bearing cytosine methylation-dependent mutations based on Gene Ontology term classification using KOBAS2.0 (Xie et al., 2011). We observed a significant enrichment of genes belonging to GO term GO:0000156 (Fisher’s exact test, *P-value* = 0.04), which includes genes that act as response regulators to bacterial two component systems. These included quorum sensing response regulator LuxO; LuxN; cytochrome c oxidase; methyl accepting chemotaxis protein and PTS system - phosphoenolpyruvate phosphotransferase system among others. Next we found that 36% (Fisher’s exact test, *P-value* = 0.028) of the genes with cytosine methylation-dependent mutations were membrane associated genes which included membrane transporters and symporters and even a surface antigen which showed a non-synonymous C → T transition. An extensive literature survey revealed many of these genes with C → T and G → A transitions at methylated cytosines are implicated in virulence of the *V. cholerae* strains **Table 2**

**Table 2.**
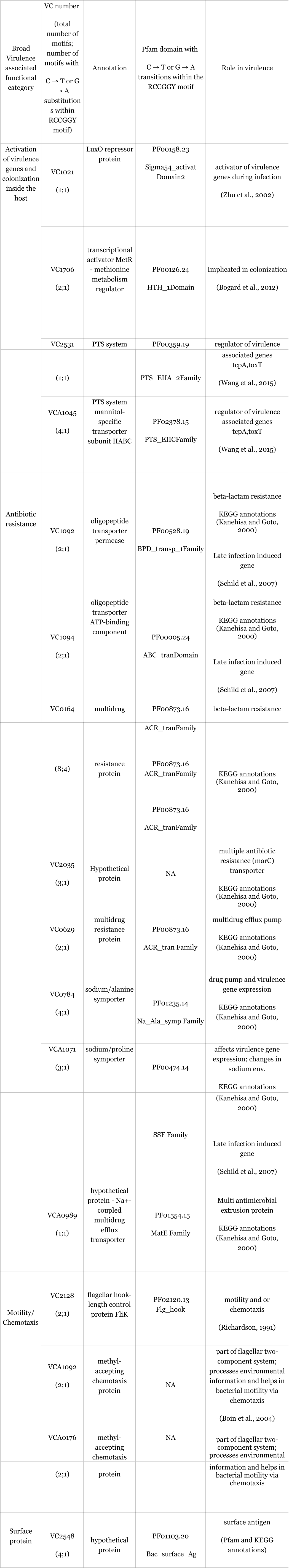

## Discussion

Epigenetic variation is widely studied in the context of its gene regulatory capacities mostly in eukaryotes and recently in bacteria (Walsh and Xu, 2006). Whereas the field of eukaryotic epigenomics is dominated by 5mC at CpG islands (Deaton and Bird, 2011), in bacteria there is increasing evidence for the abundance of epigenetic variations like 6mA, N4C and 5mC thanks to the advent of techniques like SMRT and bisulfite sequencing (Blow et al., 2016). Traditionally epigenetic variation in bacteria was known in the context of bacterial RM systems, the role of adenine methylation in core cellular processes including the cell cycle (Reisenauer et al., 1999) is being increasingly recognized. However, although the high mutational capacity of methylated cytosines to undergo deamination (Coulondre et al., 1978; Ehrlich et al., 1986; Shen et al., 1994) is well documented, the consequence of cytosine methylation-dependent mutations on bacterial genomes is less represented. Here, using genome sequence data from bacteria with known orphan cytosine methyltransferases we show the potential mutational capacity of methylated cytosines.

Previous literature and our analysis suggests that bacteria might be using motif avoidance as a strategy to overcome cytosine methylation-dependent mutations. This was true for not only *Vibrio cholerae* but for over half of the bacteria in our collection of 300 bacteria (**Figure S4**). Further, bacteria without VSR (very short patch repair protein) showed under-representation of cytosine methyltransferase target motifs as compared to the ones with VSR (**Figure S5**), implicating that the strategy of motif avoidance might be employed by many other bacterial genomes. But the motif avoidance strategy does not completely avoid the risk nor the cost associated with cytosine methylation dependent mutations, specifically since bacteria without VSR show high percentage of potential non-synonymous substitutions at methylated cytosines as compared to bacteria with VSR (**Figure S6**). Further, bacteria differ not only in the abundance of the methylcytosine target motifs, but also in terms of their methylation profiles *in-vivo*. For example in *E. coli* all the CCWGG target motifs are methylated by stationary phase (Kahramanoglou et al., 2012), but in *V. cholerae* all the RCCGGY target motifs are methylated by exponential phase itself (Chao et al., 2015). This increases the risk of C → T transitions in *V. cholerae* since (a) DNA polymerase makes errors in correctly recognizing methylated cytosines (Shen et al., 1992) (b) 5-methylcytosines on single stranded DNA at 37^o^C (temperature *V. cholerae* encounters inside human host) are more prone to deamination (Ehrlich et al., 1986) and (c) if deamination at methylated cytosines occurs on strand separation mis-match repair might fail to revert C → T and G → A transitions.

Point mutations like C → T and G → A transitions could be beneficial, detrimental or neutral to the survival of a cell. Non-synonymous substitutions in particular, by changing the proteins sequence can affect protein function (DePristo et al., 2005; Pakula and Sauer, 1989; Pál et al., 2006), and hence are mostly considered detrimental and are negatively selected. However, the occurrence of high proportions of non-synonymous methylation-associated substitutions in *V. cholerae* suggests these mutations might have been selected potentially as a result of advantages they confer to the bacterium. These mutations include those at transcriptional regulators that may play roles in virulence. Our analysis of the structure of LuxO – a quorum sensing response regulator shows that the non-synonymous C → T transition in LuxO gene changes the highly conserved glycine residue to aspartic acid in the nucleotide binding region of the protein. LuxO is crucial for activation of virulence associated genes in *V. cholerae* (Zhu et al., 2002), and a change in the nucleotide-binding region of this protein might result in a change in the expression profile. An experimental characterization of the impact of this mutation on gene expression and virulence is called for.

We also observe that membrane proteins which are one of the fast evolving proteins given their interaction partners and their role in responding to changing environmental conditions (Plotkin et al., 2004; Victor et al., 2016; Volkman, 2002), show enrichment of mutagenic cytosine methyltransferase target motifs. This suggests that cytosine methylation-dependent mutations might contribute to the observed genetic diversity at the cellular membrane. This diversity might play an important role in not only affecting bacterial host interactions but also the bacterial response to drugs since many of the membrane proteins are also drug targets. Further, abundance of orphan cytosine methyltransferases and lack of VSR homologs in pathogenic bacteria belonging to Helicobacter and Neisseria species **(Figure S7)** highlights the importance of these model systems alongside *V. cholerae* for experimentally testing the role of cytosine methylation-dependent mutations in generating genotypic diversity with an adaptive potential.

## Acknowledgements

M.S is supported by Institute of Bioinformatics and Applied Biotechnology (IBAB) in part by a grant from the Department of Biotechnology (DBT), India (BTPR12422/MED/31/287/2014, valid November 2014 to 2017) and DBT-JRF [(DBT/JRF/BET-16/I/2016/AL/86-466)]. S.K. is supported by a fellowship from the Council for Scientific and Industrial Research, India [(09/860(0122)/2011-EMR-I)]. A.S.N.S. is funded by a Ramanujan Fellowship (SR/S2/RJN-49/2010) from the Department of Science and Technology, Government of India; Indo–French Centre for the Promotion of Advanced Research (IFCPAR/CEFIPRA) grant (5103-3) and National Centre for Biological Sciences (NCBS) core budget.

